# Effectiveness of Cold Atmospheric Plasma on Staphylococcus Aureus Colonies on Living Animal Tissue Surface

**DOI:** 10.64898/2026.04.29.721726

**Authors:** Fatemeh Shakeri, Hassan Mehdian, Mahdiyeh Bakhtiyari-Ramezani, Elaheh Amini, Kamal Hajisharifi

## Abstract

*Staphylococcus aureus* (*S. aureus*) is the most common pathogen associated with skin infections worldwide. Significant efforts have been made to identify and develop innovative therapeutic strategies against *S. aureus* as alternatives to conventional antibiotics. Physical plasma has a broad range of potential uses, with non-destructive disinfection being one of its earliest applications. Although the literature emphasizes the antibacterial properties of cold atmospheric plasma (CAP), the effect of plasma on *S. aureus* on damaged skin susceptible to *S. aureus* invasion through the itch-scratch cycle has not been studied to date. Thus, we examined the effectiveness of CAP treatment on *S. aureus* bacteria in atopic dermatitis lesions using floating electrode dielectric barrier discharge devices, as well as helium and argon plasma jets. Heat distribution on the skin target, ultraviolet C radiation, and ozone generation of plasma jets for the operator of plasma sources were evaluated. Microbial tests confirmed the presence of *S. aureus* on the lesions of the groups before treatment. The groups exposed to plasma treatment showed a notable reduction in bacterial population compared to the model group (p<0.05). Furthermore, our investigation indicated that plasma treatment reduced pruritus behavior. The findings suggest that cold atmospheric plasma treatment may potentially target skin infections caused by *S. aureus* in addition to conventional therapies.

## 1- Introduction

*Staphylococcus aureus* (*S. aureus*) is a Gram-positive, spherical bacterium belonging to the family Micrococcaceae, commonly found in both human and animal environments. Regardless of age, climate, or geographical location, *S. aureus* is the most frequently linked pathogen to skin infections worldwide(1), resulting in more severe treatment failures due to antibiotic resistance. *S. aureus* causes various infections through its growth, spread in tissues, and release of several extracellular substances(2, 3). Inflammation of the skin is exacerbated by infections and toxins produced by *S. aureus*(4). Recent studies have highlighted the role of S. aureus in the development of atopic dermatitis (AD)(5), and there is evidence that the skin microbiota is more susceptible to *S. aureus* exposure during flares of the condition (6). Therefore, new and effective antimicrobial methods are always in demand for skin and skin diseases. One of these promising and growing technologies is cold plasma.

Plasma, described as the fourth state of matter, is an ionized gas that consists of free electrons and ions whose collective behavior is dominated by long-range electric and magnetic fields. Plasma is generated under different temperatures and gas pressures in a laboratory. Meanwhile, plasma with a gas temperature significantly lower than the electron temperature is classified as a non-equilibrium plasma or cold plasma(7). Nowadays, cold plasma at low pressure is commercially used for the sterilization of medical tools(8). However, vacuum limitations for sterilizing living tissue led to the proposal of techniques to generate cold plasma at atmospheric pressure (CAP), and plasma medicine science gained immense scientific interest, prompting many researchers to conduct further studies. The main components of CAP are charged particles, electric field, oxygen and nitrogen species, ultraviolet (UV), and vacuum ultraviolet radiation(9).

Various configurations of cold plasma sources can be used for sterilizing tissue. The floating electrode dielectric barrier discharge (FE-DBD) used the tissue as one of its electrodes. A plasma jet or pen produces plasma between electrodes, and the components are transported to the target, making it an excellent prospect for use in biological and medical applications(9-13). Studies demonstrated the significant antibacterial properties of cold plasmas, especially on nonliving surfaces. For example, Nicol et al. (14) utilized a plasma jet with helium/oxygen admixture flows to inhibit the growth of *S. aureus* and *Escherichia coli* bacteria on solid and porous surfaces. Dijksteel et al. (15) investigated the safety and efficacy of a flexible surface DBD device for treating infected wounds. The generated plasma was effective against *pseudomonas aeruginosa* in vitro and in vivo.

Limited research has focused on the significance of interactions and the diverse behaviors of bacteria on live skin compared to culture medium. Daeschlein et al.(16) investigated the in vivo decontamination potential of plasma jet and DBD on human fingertips. In addition, other reports have demonstrated that plasma jet devices can effectively inactivate both multidrug-resistant and non-resistant bacterial pathogens in chronic wounds(17, 18). Even though the studies mentioned above indicate that CAP has antibacterial effects(16, 19-22), no research has been conducted on the impact of various plasmas on *S. aureus* of damaged skin susceptible to invasion through the itch-scratch cycle. The itch-scratch cycle causes continuous mechanical damage to the skin and colonization of pathogens. Therefore, stopping the repetitive cycles of scratching and itching and restoring the damaged skin barrier may be an effective treatment strategy for AD. Research indicated a correlation between skin colonization by *S. aureus* and the intensity of pruritus, suggesting *S. aureus* may play a role in the aggravation of itching associated with AD (23, 24).

Here, we aim to investigate the effect of floating electrode dielectric barrier discharge, helium (He), and argon (Ar) plasma jets treatment on the colonization of *Staphylococcus aureus* and pruritus behavior on AD-like lesions induced by 1-chloro-2,4-dinitrobenzene (DNCB) in the BALB/c mice model. Since Tacrolimus (TAC) ointment is a calcineurin inhibitor that is effective for the treatment of atopic dermatitis. The effect of TAC was also investigated on the *S. aureus* colonization and pruritus behavior on AD-like lesions. Moreover, we assessed plasma sources regarding heat distribution on the skin target, UVC radiation, and ozone (O_3_) generation, which are probably involved in plasma disinfection properties, for the safety evaluation of plasma sources on the target and operator. Due to differences in the configuration of plasma sources and the ionization potential of gases, resulting in unique concentrations and compositions of reactive species, a comparison of sources is beyond the scope of this study. To provide an outline of the paper, we describe the experimental setup, including plasma sources, measurement methods for plasma, and microbiological characteristics. The analyzed results are then discussed.

## 2- Materials and methods

### 2-1- Physical characteristics of plasma sources

#### Experimental setup

Figure 1(a) depicts the experimental setup of a plasma source, including a power supply, an electrical measurement system, a gas supply (only for jets), and an optical detection system. Figure 1 (b,c,d) represents the schematic plasma jets and FE-DBD devices. The helium plasma jet is based on the DBDjet configuration (25) (Figure 1(b)) and was driven by a high-voltage AC power supply operating at a frequency of 12 kHz and a voltage of 7 kV. The helium gas flow was kept at 4 slm. Argon plasma jet (Figure 1 (c)) is equipped with a stainless steel needle-shaped electrode (0.1 mm) located at the center of the alumina tube, which is connected to a power source (100 kHz pulse wave modulated into a 300 Hz pulse with a 35% duty cycle). The gas flow rate was adjusted to 5 slm. The helium and argon gases, of 99.998% purity, were fed to jet devices to minimize toxic gas emissions. The distance between the nozzle and the skin surface of the mice was 10-15mm. A homemade FE-DBD device consists of a copper rod as a central electrode and a Pyrex tube as a dielectric barrier. Figure 1(d) shows that the skin serves as the second electrode of the FE-DBD device. The dielectric limits the electrical current and avoids arcing. The discharge ignited at a voltage of 14 kV at 25 kHz when the distance between the high-voltage electrode and the surface was less than 3 mm, as shown in Figure 1(d).

**Figure 1.**
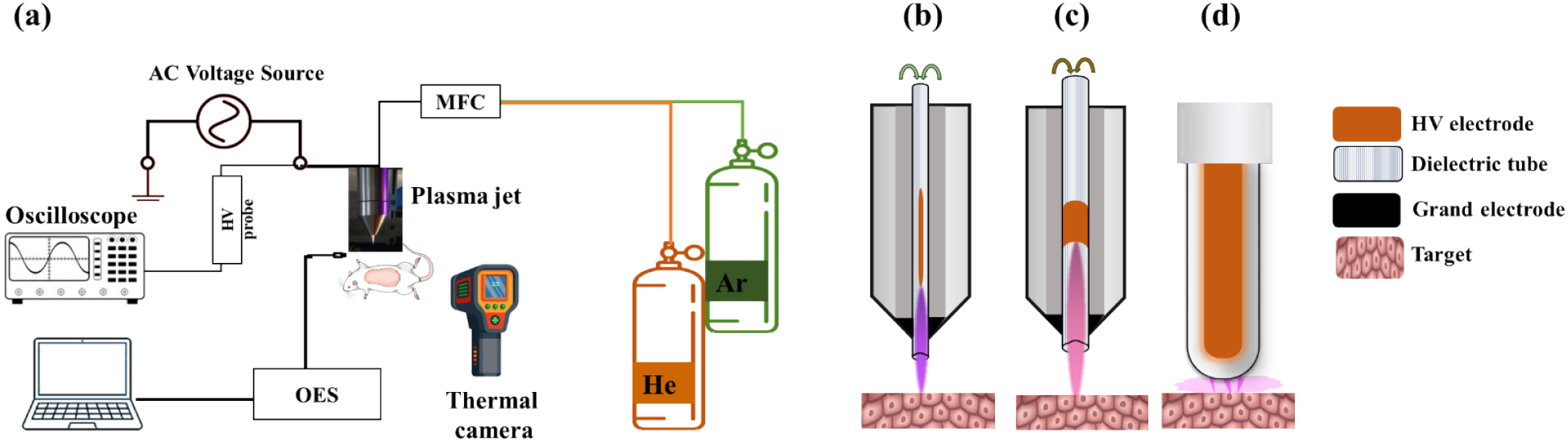
Schematic of the (a) experimental setup, (b) helium plasma jet, (c) argon plasma jet, (d) FE-DBD device

#### Optical emission spectroscopy

Optical emission spectroscopy (OES) was used to determine the ionization stages of the ions or molecules in the plasma by OES spectroscopy (Optical Physics Technologists Company, Iran). The wavelength range of the spectrometer was 200-900 nm. The spectra were recorded at an axial position, 10 mm from the nozzle outlet, with an acquisition time of 1 s. Before data acquisition, the jets were switched on for approximately 30 s until no fluctuations in the spectra were visible. The spectrum was analyzed by software (Version 1.14). In this study, the OES of the FE-DBD device was not measured. The spectrum was averaged over 20 scans to remove fluctuations.

### 2-2- Safety evaluation setup for plasma therapy

#### Infrared thermal imaging

Infrared thermal imaging is a non-contact, non-destructive technique that allows for the mapping of an object’s temperature. The mouse skin temperature was monitored immediately after CAP treatment with an infrared thermal imaging camera model 868 (Testo, UK), and the images were analyzed using IR software (Version: v4.8).

#### UV radiation measurement

The UV radiation intensity generated by the CAP sources was measured using a calibrated UV radiation meter UVC-254A (Lutron, Taiwan). The sensor provides a spectral response of 200 to 280. The UV sensor was utilized to measure the UVC intensity of CAP sources for specific parameters on the device.

#### Measurement of ozone concentration

The accumulation of ozone concentration in a glass box (0.5 m × 0.5 m × 0.5 m) is determined for a 3-minute plasma treatment using an ozone analyzer (Horiba APOA-370, Japan). The analyzer was set to a range of 0-0.1 parts per million (ppm), providing a measurement accuracy of ±0.001 ppm within our measured concentrations. The ozone concentration of FE-DBD could not be measured due to the device’s construction.

### 2-3- Microbiological assay of plasma sources

#### Animals

A total of 18 female BALB/c mice (6–8 weeks old, 20±2 g) were purchased from the Experimental Animal Center of Pasteur Institute of Iran (Tehran, Iran). All animal experiments were undertaken according to the guidelines of the Animal Care Committee of Kharazmi University (approval number IR.KHU.REC.1402.069). All mice were housed in groups of three per cage in a modeled environment, with a constant temperature (22 ± 2°C), humidity (45-55%), and a 12-h light-and-dark cycle. Food and water were freely available. The mice were randomly divided into six groups of three animals each, using a computer-generated randomization list, as follows: The non-intervention group (normal; G1), the DNCB-sensitized group (model; G2), the helium plasma-treated group (G3), the argon plasma-treated group (G4), the air plasma treated-group (G5) and the Tacrolimus ointment (Protopic 0.1%) treated-group (G6).

#### AD Model induction protocol, treatment, and sampling

The atopic dermatitis model was selected to investigate the effect of plasma on removing *S. aureus* bacteria from living skin. After 1 week of acclimatization of female BALB/c mice, the hairs of the entire back of mice in all groups were removed from anesthetized mice using an electric shaver and hair removal creams. Then, mice were sensitized with 100 μL of 1% DNCB (Sigma–Aldrich, St. Louis, MO, USA) in acetone: olive oil (4:1) on days 1 and 2. On day 3, 120 μL of 2% DNCB in acetone: olive oil (4:1) solution was applied to dorsal skin. Mice were then received 100μL of 0.5% DNCB twice to maintain inflammation (26-28). The treatment area (∼2cm^2^) was defined on the skin using a template and subsequently tracked throughout the study. As shown in Figure 2, clinical signs of dermatitis, including dryness/erosion, edema, erythema/bleeding, and excoriation, were clearly observed on day 7, prior to treatment. After confirming the AD model, all treatments were performed eight times on days 7-14. CAP sources were moved uniformly across the entire treatment area for a total of 2 minutes (1 min/cm^2^), while 100 mg/mouse of 0.1% TAC ointment was used on the AD-like skin. Briefly, microbial sampling was performed on days 7, 10, and 14 using sterile cotton swabs pre-wetted in sterile normal saline. The swabs were gently rotated five times over a 1 cm^2^ dorsal skin lesion to ensure uniform bacterial collection. Each swab was then transferred to a sterile tube containing 1 mL of sterile normal saline.

**Figure 2.**
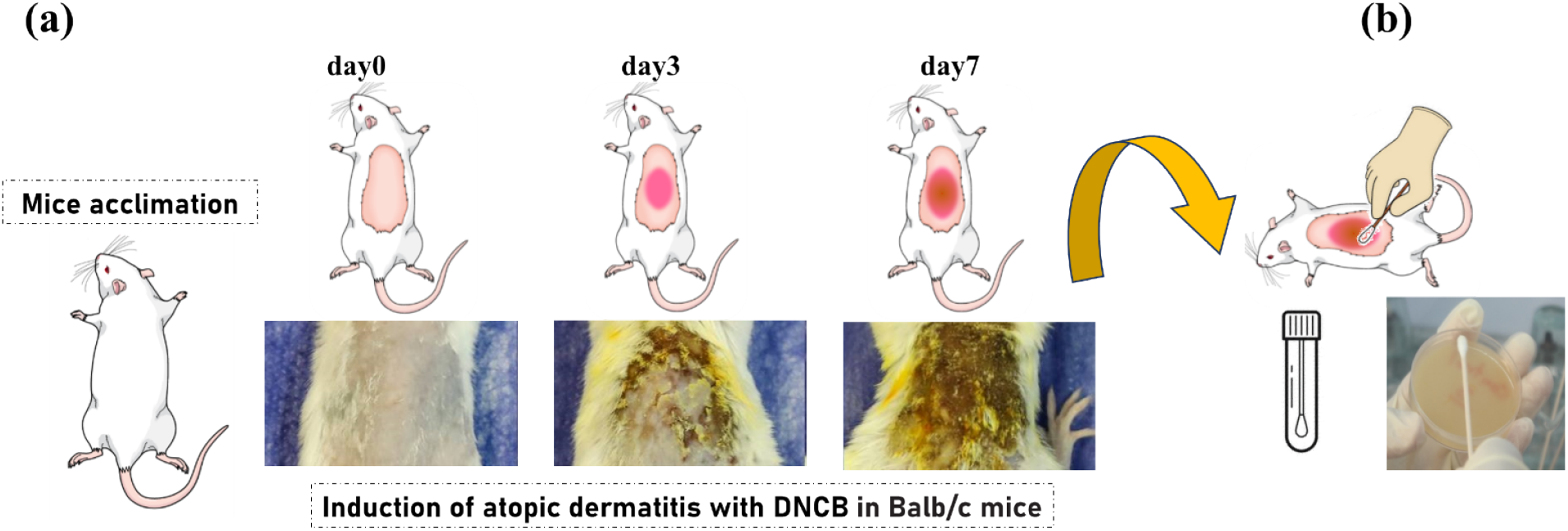
(a) Temporal changes in AD mouse model induction, (b) Sampling and microbiological assay

#### Baird Parker Agar

Baird Parker Agar (Merck, Germany) was used to isolate and enumerate *S. aureus*. The medium also allows the differentiation of coagulase-positive strains(29, 30). The desired quantity of the medium was suspended in 950 mL of purified water, then heated with frequent agitation and boiled to dissolve the medium completely. Afterward, it was autoclaved at 121°C for 15 minutes and cooled to 45-50°C per the manufacturer’s instructions. The medium was then poured into sterile petri plates under sterile conditions. The 0.1ml of normal saline containing diluted bacteria was spread on the agar surface until dry. After 24-48 hours of incubation, colonies of *S. aureus* appeared. The experiment was conducted in triplicate. Each 1 cm^2^ surface sample was eluted in 1 mL of sterile saline; therefore, the measured CFU/mL values were considered equivalent to CFU/cm^2^.

#### Mannitol Salt Agar

Mannitol Salt Agar (MSA) selects for Staphylococcus species that can tolerate high salt concentrations. To prepare MSA (Merck, Germany), 108.0 g was suspended in 1000 mL of distilled water and boiled to ensure complete dissolution. It was sterilized by autoclaving at 121°C for 15 minutes as per the manufacturer’s instructions. The medium was then poured into sterile petri plates under sterile conditions. The swab samples were streaked onto MSA and incubated at 37°C for 24 hours to collect bacterial isolates that were able to grow on this medium. Colonies were identified by colony morphology and the presence or absence of mannitol fermentation on MSA. Single colonies were tested using deoxyribonuclease and tube coagulase tests, as well as growth on MSA(31).

#### Deoxyribonuclease and Coagulase tests

A deoxyribonuclease (DNase) assay was performed to presumptively identify *S. aureus* and differentiate it from other Staphylococcal species. The medium was added to 1000ml of pure distilled water. The dissolved medium was autoclaved at 121°C for 15 minutes and then cooled to about 40-45°C. The medium was then poured into sterile Petri plates under sterile conditions. The colonies were cultured in the presence of DNase and incubated for 24-72 hours at 37°C. A zone of visibility around the colony indicates a positive test outcome. A tube coagulase test (TCT) was also performed on the colonies. The 0.5ml of Rabbit plasma with Ethylene diamine tetraacetic acid (Baharafshan, Iran) was transferred into the test tube. The isolated colony was collected using the sterile loop. After emulsifying, the tube was placed in the incubator. The samples were checked 1-4 hours for evidence of a clot. The test continued with an overnight incubation for any clot formation, considered a positive result, and for a negative result.

#### Eosin Methylene Blue agar

Eosin Methylene Blue (EMB) agar (Merck, Germany) effectively suppresses the growth of various Gram-positive bacteria, and the absence of isolated bacterial growth indicates that accurate sampling and proper selection of bacterial populations have been achieved. To prepare EMB, 36 g was suspended in 1000 mL of distilled water to create a uniform suspension, and then sterilized by autoclaving at 121°C for 15 minutes. The suspension was cooled to 45-50°C, as per the manufacturer’s instructions. The medium was then poured into sterile petri plates under sterile conditions.

### 2-4- Assessment of pruritus behavior

A digital video camera (Canon 5D Mark IV, Japan) was used to record the pruritus for 10 minutes per mouse. Back pruritus was counted, while other sites, such as the ears, were disregarded. The continuous scratching of the back, which persisted for more than three seconds, was scored as two distinct scratching events. In other words, for longer durations, it was counted in intervals of three seconds. The data were collected in triplicate (three mice per group), resulting in a total of 30 minutes of observation per group. One blinded observer gathered the data to prevent variations in the definition of an itch among observers(32, 33).

### 2-5- Statistical analysis

Sample size for each group was determined based on a power analysis, using effect sizes estimated from pilot data, to ensure at least 80% statistical power. After testing normality, results were evaluated by analysis of variance, and statistical analyses were performed on three replicates of data obtained from each treatment. Comparisons among multiple groups were performed using a two-way ANOVA repeated measurement test followed by Duncan’s multiple comparison post hoc test. The statistical significance level was set at p<0.01 and p<0.001. All data analysis was performed using SPSS 27.0, TIBCO Statistica™ 14.0.0, and PRISM 9.5.1 (GraphPad software) and G*power.

## 3- Results

### 3-1- Optical emission spectroscopy characterization of CAP devices

Figure 3 represents the spectral lines of the He and Ar plasma jets emitted in the Ultraviolet-visible regions in the described experimental setup. The propagation of the plasma jet into the air results in interactions with molecules, which subsequently lead to the formation of reactive species. As shown in Figure 3 (a), essential peaks in the emission of the helium plasma jet are attributed to atomic He (587 nm, 667 nm, 706 nm, and 728 nm) and atomic oxygen (777nm) as products of ambient O_2_ and H_2_O dissociation, OH radical (306-312nm), NO (λ = 230-260 nm)(34, 35), excited molecular nitrogen (357nm) and ionized nitrogen (391nm) as helium metastable presence indicator. It is observed that the spectrum is dominated by the N2 second positive, and the spectral lines of atomic helium indicate the presence of numerous energetic species in the helium plasma jet (36-38). Figure 3(b) represents the results of the reactive species intensity of the cold argon plasma jet, where the species were identified in the spectrum. The lines of argon atoms are apparent in the 680-850nm spectral range. Moreover, the emission spectrum demonstrated the presence of hydroxyl radicals, excited nitrogen, and excited oxygen in the argon plasma(38). In the present setup condition, the emission intensity of OH was higher in the argon than in the helium plasma jet, and NO was not detected in the argon plasma jet. The OES results revealed that both the helium plasma and the argon plasma jet produce excited nitrogen and hydroxyl radical species. The helium plasma was more efficient in exciting N_2_ and forming NO, while the argon plasma appeared superior in generating OH radicals.

**Figure 3:**
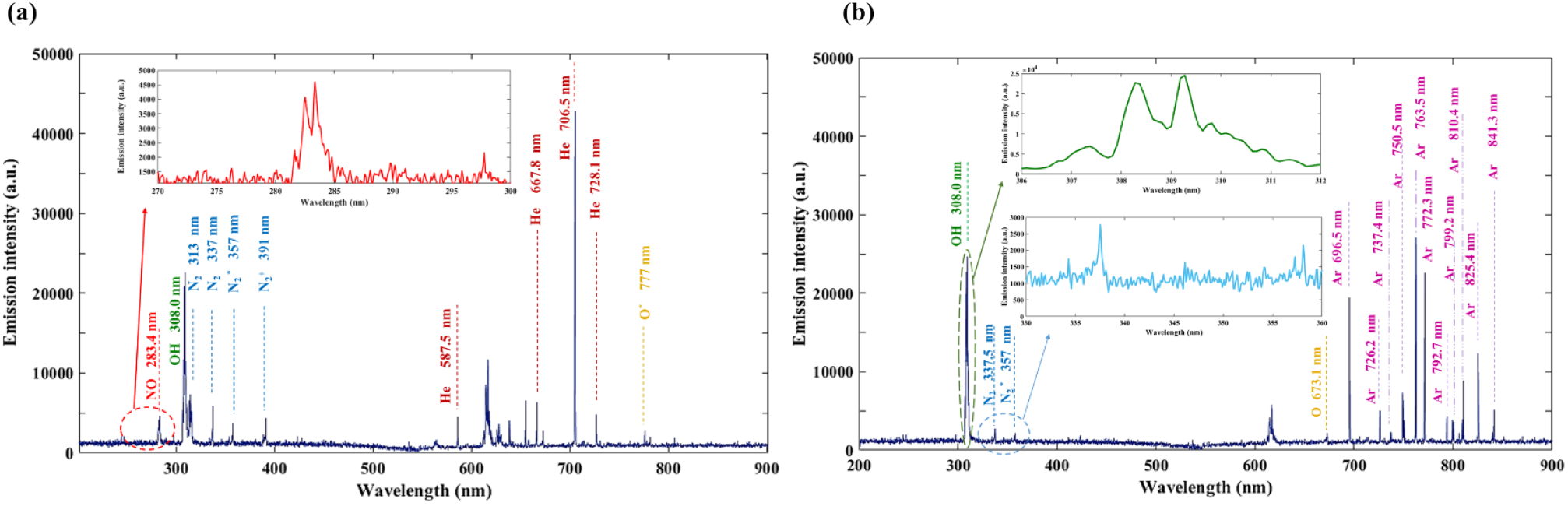
Representative optical emission spectra of (a) helium plasma and (b) argon plasma jets averaged over 20 scans at experimental conditions

### 3-2- Safety measurement results for plasma therapy

The temperature of the AD-like skin was approximately 30 °C. During plasma treatment using jets and FE-DBD, the temperature rises to a maximum of 35 °C, as shown in Figure 4. The temperature distribution of jets and FE-DBD is different, as shown in the circle region in Figure 4. The jets increase the temperature of the skin point under plasma treatment, while FE-DBD has a broad region. This is remarkable for point-shaped plasma sources. On the other hand, various wavelengths of UV radiation (UVC: 100–280 nm, UVB: 280– 315 nm, and UVA: 315–400 nm) can be obtained from plasma sources. To characterize the UVC, the intensity of the device during operation was measured. The intensity at the mentioned setup was measured to be approximately 0.2 ± 0.1 µW/cm^2^ and 0 ± 0.1 µW/cm^2^ for the He and Ar plasma jets, respectively. The amount of VUV radiation (120-150nm) emitted from the plasma devices was not quantified in this study. The second product of plasma sources is ozone, a highly reactive gas that acts as a potent oxidizing agent. The ozone concentrations were measured to ensure the operator and the animal’s safe use of plasma devices. The initial ozone concentration was 0.007 ± 0.001 ppm. The mean ozone concentration (measured over 3 minutes) was 0.017±0.001 ppm and 0.012±0.001 ppm for He and Ar plasma jets, respectively.

**Figure 4:**
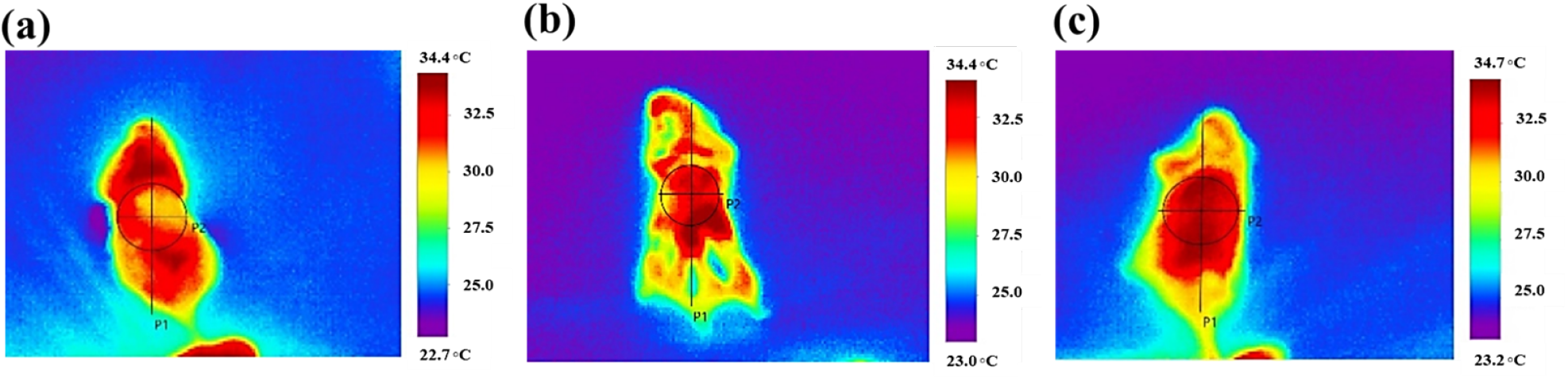
Thermal distribution of animal skin a few seconds after plasma treatment by (a) helium plasma jet, (b) argon plasma jet, and (c) floating electrode dielectric barrier discharge devices

### 3-3- CAP effects on *S. aureus* colonization on AD-like skin

The identification of *S. aureus* on the AD-like lesion was confirmed by combination tests (MSA/DNase/TCT). The results showed clear zones around the bacterial colonies (DNase-positive), clot formation in the tube (TCT-positive), and no bacterial growth on EMB. Figure 5 shows the changes in the logarithm of the *S. aureus* population in different experimental groups and on various days. As shown in Figure 5, the logarithm of the bacterial population was the lowest for all experimental days in the normal group (G1) and the highest for all experimental days in the model group (G2). All experimental groups exhibited the highest logarithmic values of bacterial populations prior to treatment, with no statistically significant difference (P > 0.05). There were no statistically significant changes in bacterial load after TAC ointments were administered on the AD-lesion in the G6 group on days 10 and 14 compared to before treatment. On the other hand, AD lesions treated with CAP in the G3, G4, and G5 groups recorded a low bacterial count compared to the initial count on day 7. The group subjected to helium plasma treatment (G3) showed a notable reduction in bacterial population on day 10 compared to the model group (G2) and TAC-treated group (G6). This downward trend in bacterial counts persisted significantly through day 14 of the treatment. The group exposed to argon plasma treatment (G4) demonstrated a significant reduction in bacterial population on day 10 following four consecutive treatments, in comparison to day 7 and both the model and ointment groups. However, an increase in bacterial population was observed on day 14 for this group, although this change was not statistically significant when compared to day 10. The air plasma-treated group (G5) showed a reduction in bacterial populations on day 10 compared to day 7, in both the model and TAC groups. However, the bacterial reduction rate was slower on day 14 (as seen in Figure 5).

**Figure 5:**
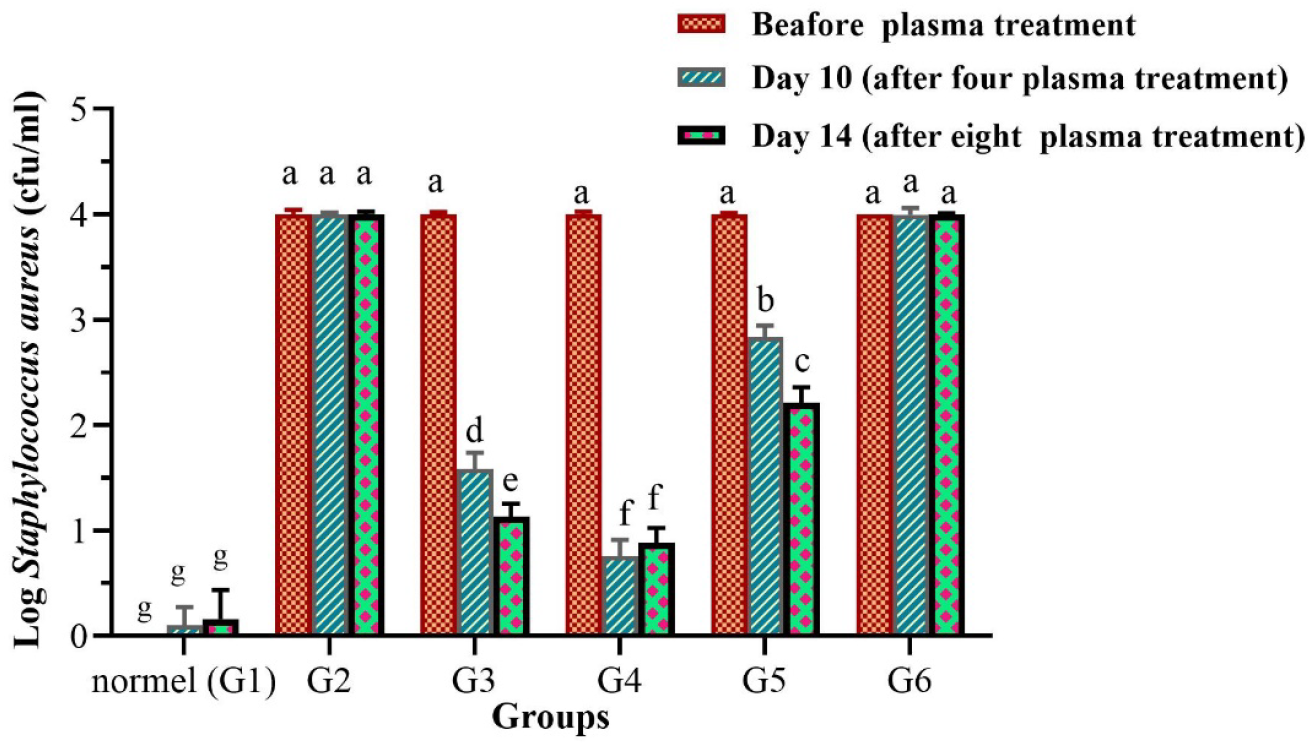
Effects of CAP sources and TAC treatment on S. aureus colonization in AD-like mice (CFU/ml = CFU/cm^2^). Logarithmic values of S. aureus of dorsal skin in each experimental group (G1–G6) on days 7, 10, and 14. Data represent mean ± SD; n = 3 mice per group. G1(normal), G2 (model), G3 (helium plasma-treated group), G4 (argon plasma-treated group), G5 (air plasma-treated group), and G6 (TAC-treated group). * Means followed by the same letter are not significantly different (p<0.05).

The progressive bacterial reduction observed in G3 and G5 on day 14 was more pronounced than on day 10, indicating the time-dependent antibacterial effect. A statistically significant reduction in bacterial population was noted in the G3 and G4 groups compared to the measurements taken on the fourth day of plasma treatment. This suggests a progressive decline in the bacterial population resulting from the use of helium and air cold plasma. Changes in the logarithmic values and percentages of bacterial populations in the G3, G4, and G5 groups on days 10 and 14 following CAP treatment in comparison to the G2 group are shown in Table 1. In other words, the % represents the relative reduction in bacterial load in comparison to the model group on the respective days.

**Table 1.** log S. aureus population decrease observed in each experimental group on days 10 and 14 compared to the model group (G2).

|  | Groups | Logarithmic |
| --- | --- | --- |
| Day 10 (After four times plasma treatment) | G3 | 2.41 |
|  | G4 | 3.24 |
|  | G5 | 1.16 |
| Day 14 (After eight times plasma treatment) | G3 | 2.86 |
|  | G4 | 3.11 |
|  | G5 | 1.78 |

### 3-4- CAP effects on pruritus of an AD mouse model

Figure 6 shows the changes in pruritus behavior across different experimental groups and on various days. The evaluation results before plasma treatment indicated that the pruritus was the most severe across all experimental groups, except the normal group (G1). In the normal group, pruritus was the lowest. The observation of pruritus behavior in mice within the normal group over the periods of 10 and 14 days indicated a minor increase; however, this change was not statistically significant during the observation period (P > 0.05). There were no substantial changes in pruritus behavior in the model group on days 10 and 14 after the induction of AD. The persistence of pruritus in the model group over the observation period reflects the effect of repeated DNCB sensitization and challenge. CAP treatment reduced the pruritus behavior in the G3, G4, and G5 groups compared to the model group, with significant improvement observed in the argon plasma-treated group (Figure 6). As mentioned in Table 2, the decrease is statistically comparable to that of the model group. In general, after four subsequent plasma treatments, a significant reduction in pruritus was observed in both the helium and air plasma groups compared to the pre-treatment levels. The plasma treatment extended to the 14th day of AD induction (eight treatments) contributed to the reduction of pruritus behavior. The pruritus behavior assessment of the argon plasma-treated group observed was statistically the same on day 10 as on day 14 (p>0.05). Following evaluation, the helium and air plasma-treated groups also showed a continuous decrease in itch on the days (Figure 6). The application of TAC ointment in the G6 group resulted in a significant reduction in pruritus, recorded on days 10 and 14 after AD induction, compared to measurements before treatment. No significant changes were recorded in the pruritus intensity during the four-day observation period. The reduction of pruritus in this group was much lower compared to the G3, G4, and G5 groups. The percentage of changes in pruritus behavior in the G3, G4, and G5 groups compared to the G2 group on the 10 and 14 days of AD induction is summarized in Table 2.

**Table 2.** The pruritus decrease observed in each experimental group on days 10 and 14 compared to the model group (G2).

|  | Groups | % |
| --- | --- | --- |
| Day 10 (After four times plasma treatment) | G3 | 49.33 |
|  | G4 | 77.33 |
|  | G5 | 41.33 |
| Day 14 (After eight times plasma treatment) | G3 | 81.18 |
|  | G4 | 88.23 |
|  | G5 | 61.18 |

**Figure 6:**
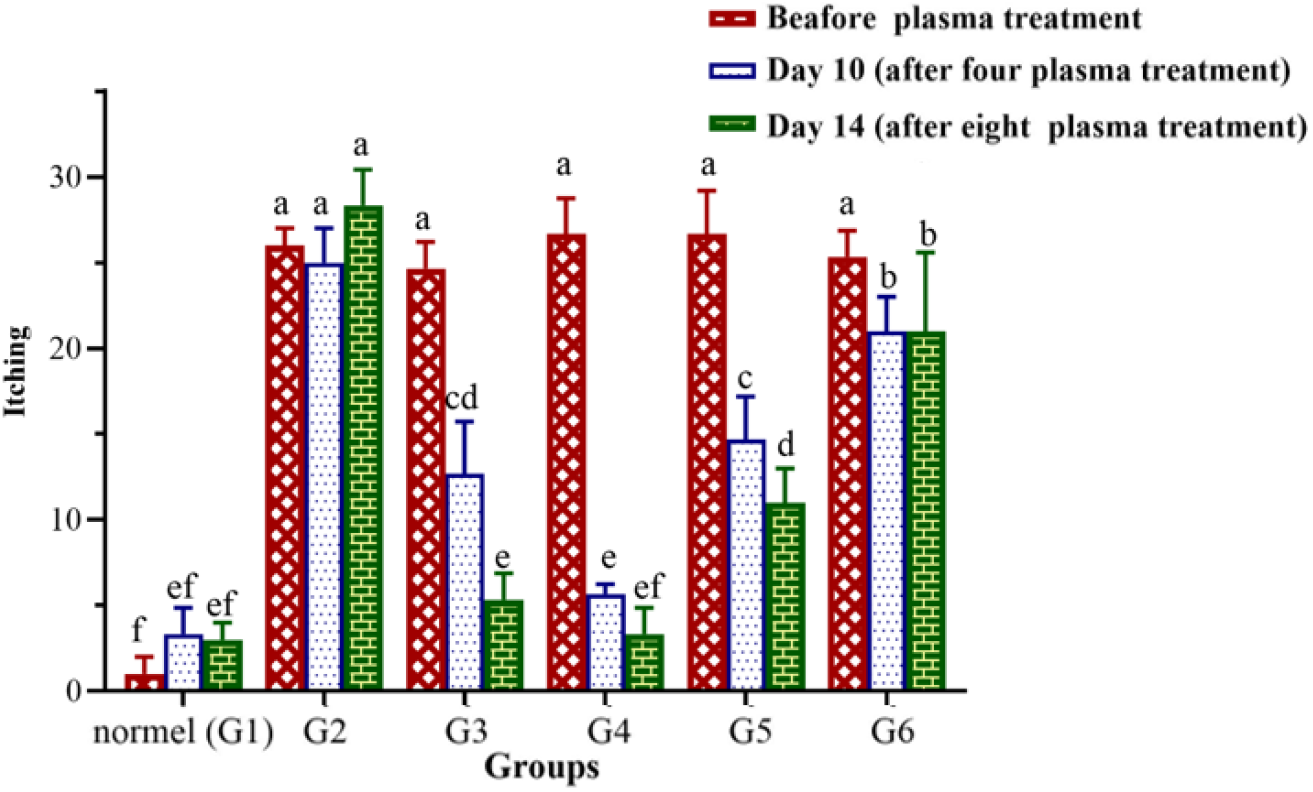
Effects of CAP sources and TAC treatment on pruritus behavior in AD-like mice. (a) itching scores of each experimental group (G1–G6) on days 7, 10, and 14. Data represent mean ± SD; n = 3 mice per group. G1(normal), G2 (model), G3 (helium plasma-treated group), G4 (argon plasma-treated group), G5 (air plasma-treated group), and G6 (TAC-treated group). * Means followed by the same letter are not significantly different (p<0.05).

## 4- Discussion

Skin colonization of *S. aureus* is seen in approximately 90% lesional skin and 55% to 75% non-lesional skin(39). *S. aureus* is known to colonize the skin of AD patients and is the most common organism responsible for wound infections after surgery(40, 41). It was found that *S. aureus* releases proteases, enzymes, and toxins that destroy cells by causing damage to the cell basement membrane and can exacerbate barrier dysfunction.(42). Over the past several decades, dealing with multidrug-resistant strains has been challenging for treating staph infections due to the mounting number of drug-resistant strains(43). In response to the increasing mortality and medical costs of drug-resistant bacteria, new treatment options are becoming more prevalent. Hence, innovative therapeutic approaches are needed to eliminate *S. aureus*. Using ultraviolet light, hydrogen peroxide, and ozone disinfection methods can effectively kill S. aureus, thereby preventing it from infecting the human body (44-46).

CAP is an alternative disinfection method that can eliminate pathogens, such as *Escherichia coli* and *Enterococcus faecalis* (47, 48). This method is proper for disinfecting wounds after medical surgery to prevent the aforementioned problem of wound infection by *S. aureus* (49). In the present study, we examined the impact of atmospheric pressure cold plasma, generated using different gases and configurations, on *S. aureus* colonization in an animal model of AD. Evidently, the first step in applying plasma devices is ensuring safety for both the patient and the operator. Although a comprehensive safety evaluation of CAP devices is beyond the scope, we focused on certain parameters, such as UVC radiation, ozone generation, and heat distribution, that may be involved in plasma disinfection and potentially harmful to the operator. A plasma temperature of below 40°C is recommended for treating living surfaces. The temperature at the end of the capillary nozzle drops with an increased distance (50, 51). In this way, IR imaging of the skin and a touch of the plasma plume indicated low temperature and safe operation. Moreover, treatments with the FE-DBD device on animal skin revealed no dramatic changes in temperature. The observed minor change was far below the dangerous threshold (less than 40 °C), which agrees with temperature measurements on similar plasma devices (52, 53). Bacterial cells can be eliminated by heating, especially S. aureus, at 80 °C for 20 minutes. The quantity of bacteria destroyed varies depending on the heat intensity and duration(54). Heat is not responsible for the observed sterilization effect, but it is likely a contributor to it through interaction with other plasma components.

The UVC intensity of CAP devices depends on device-specific settings, including distance, power, gas flow, angle, and exposure time. The UVC intensity decreases with the distance from the plasma as a 1/r^2^ function, and the highest intensity is related to the longitudinal axis of the plasma(55). Exposure to UVC in high doses can result in direct damage to nucleic acids and proteins, leading to genetic mutations or cell death (56). The germicidal strength of UV light, particularly UVC, has been established for over 100 years(56). The impact of UV radiation on the disinfection efficiency of plasma is determined by its emission in the UVC range(57). By altering the chemical structure of DNA, UVC damages cells and thus inactivates microorganisms(58). Furthermore, UVC reaches the upper layer of the epidermis(59). In this regard, our OES spectra result and the power radiation measurement of CAP devices revealed low UVC intensity in our experiment. This result can be attributed to the absorption of UVC emitted by CAP devices by oxygen molecules in the air. Thus, the UVC plays no significant role in the plasma device sterilization process under these experimental conditions. These results align with reference (57).

Besides UV radiation, the ozone production of plasma sources poses a challenge to medical treatment. Ozone in the air can affect human lungs if its concentration is above safe levels, but it does not harm the target tissue. Although O3 is beneficial for sterilization usage(46, 60, 61) and involved in chemical reactions of plasma-generated active species(9), inhalation of ozone gas can affect human lungs if its concentration in the air is above safe levels(62). The Food and Drug Administration requires the ozone output of indoor medical devices to be no more than 0.05 ppm. The Occupational Safety and Health Administration mandates that workers not be exposed to concentration levels exceeding 0.10 ppm for more than 8 hours. For a treatment time of 3 min, the threshold of 0.05 ppm is not reached. However, for repeated applications, proper ventilation, adjustment of treatment intervals, and monitoring may be required to prevent potential ozone accumulation. This consideration could be addressed in future studies.

To evaluate the effect of CAP on *S. aureus* in damaged skin susceptible to *S. aureus* invasion, DNCB, the most well-known allergen-inducing AD, was used(28, 63). Indeed, after repeated application of DNCB, the damaged skin provided an ideal environment for colonization by the desired bacterium, *S. aureus*. DNCB increased *S. aureus* colonization and induced pruritus behavior. CAP has demonstrated effectiveness in eradicating S. aureus within the unique and complex microenvironment of the skin; however, the treatment process and elimination rate may vary depending on the type of plasma used. As mentioned, helium plasma treatment of AD-like lesions resulted in a 2.41 log reduction in S. aureus load after four consecutive treatments, and this reduction increased to approximately 2.86 log after eight treatments. This trend demonstrates a significant decrease in *S. aureus* load after consecutive treatments. Argon plasma treatment reduced *S. aureus* load in AD-like lesions. However, the reduction did not significantly change between day 10 and day 14, indicating that additional consecutive treatments did not further enhance its effect. Thus, while argon plasma is effective, its time-dependent improvement is less pronounced than that of helium plasma. Although an in vitro study has exhibited that the DBD device was more effective against *S. aureus* than argon plasma jet (64), our results showed that the effect of air plasma with a DBD device is less than that of plasma jets and reached a 1.78 log reduction after eight consecutive treatments in AD-like lesions. This may be due to the fact that AD-like lesions are not smooth and involve lichenification. Hence, we suspect that the plasma jet facilitated the treatment of uneven and heterogeneous surfaces, resulting in a more effective elimination of *S. aureus*. Though a statistically significant reduction for the helium jet was observed on the final day of assessment, this can be interpreted as a result of various configurations, power supplies, and working gases involved in the chemical composition and reactivity of plasma(7).

We also examined the effect of plasma therapy on pruritus behavior as an indirect measure of *S. aureus* skin colonization. It has been demonstrated that *S. aureus* skin colonization may be a factor that exacerbates itch in AD(23, 65, 66), and restoring the skin microbiome may reduce the intensity of pruritus (67). Our results showed that argon plasma modulated the pruritus behavior and reduced it to normal after four consecutive treatments. The result is consistent with a reduction in the *S. aureus* load. The effect of helium plasma treatment was slower and took longer to return to normal pruritus levels. The air plasma treatment did not reach normal pruritus even after eight therapies. Given the mechanism by which plasma eliminates S. aureus, biological processes such as stress protein production, antioxidant production, and nitrosative stress response, as well as their effects on DNA repair, transcription, translation, and cell membrane/wall synthesis, are involved in prolonged treatment (20). Another possible mechanism is the promotion of macrophages’ ability to eliminate internalized bacteria, which could lead to the killing of both antibiotic-sensitive and antibiotic-resistant strains of S. aureus (68). In summary, plasma interaction increased cell membrane disruption, oxidative stress, and ultimately, cell death (69).

It is worth noting that TAC ointment has been approved for the treatment of atopic dermatitis. Although studies have demonstrated that topical immunosuppressive therapy of AD with tacrolimus ointment is associated with an overall decrease in *S. aureus* colonization on lesions (after 3 weeks)(70), our results showed that TAC controlled pruritus, but was not effective on bacterial colonization. Indeed, we believe that it depends on understanding the molecular mechanisms of action of the drug. On the other hand, pruritus in AD is the result of a complex interaction between multiple factors, and several itch mediators and associated receptors are responsible for itch in AD, one of which is colonization by *S. aureus* bacteria(39). Therefore, our hypothesis is that pruritus behavior was controlled in the TAC group by another molecular mechanism.

## 5- Conclusion

In the present study, we indicated that floating electrode dielectric barrier discharge, helium, and argon plasma jets treatment could effectively reduce the colonization of Staphylococcus aureus and pruritus behavior on AD-like lesions induced by DNCB. A significant log reduction of 2.86, 3.11, and 1.78 in bacterial population was observed compared to the model group after the helium, argon, and air plasma treatment. This result suggests that CAP sources may help reduce the pathogenic bacterial burden, which could contribute to restoring the microbial balance in AD-like lesions. However, further studies are required to evaluate the broader effects of CAP on the overall skin microbiota. In summary, the impact of CAP sources on alleviating pruritus behavior demonstrated an efficacy exceeding 60%. In addition, plasma jets demonstrated greater antibacterial efficacy than FE-DBD devices in treating AD skin, whose surface area is not smooth and flat. We will address the comprehensive study of plasma therapeutic effects in the future.

## Declaration

### Availability of Data and Materials

The datasets used and/or analyzed during the current study are available from the corresponding author upon reasonable request.

### Author contributions

**Fatemeh Shakeri:** Conceptualization, Project administration, Investigation, Methodology, Data curation, Formal analysis, Writing – original draft. **Hassan Mehdian:** Investigation, Methodology, Supervision, Writing – review & editing. **Mahdiyeh Bakhtiyari-Ramezani:** Conceptualization, Project administration, Investigation, Data curation, Supervision, Resources, Writing – review & editing. **Kamal Hajisharifi:** Investigation, Methodology, <u>Formal analysis,</u> Validation, Writing – review & editing. **Elaheh Amini:** Investigation, Methodology, Writing – review & editing

## Acknowledgements

The author would like to thank all those who generously helped in designing, conducting, interpreting the results, and writing the article of this study.

